# A gene for social immunity in the burying beetle *Nicrophorus vespilloides*?

**DOI:** 10.1101/020644

**Authors:** William J. Palmer, Ana Duarte, Matthew Schrader, Jonathan P Day, Rebecca Kilner, Francis M. Jiggins

## Abstract

Some group-living species exhibit social immunity, where the immune system of one individual can protect others in the group from infection. In burying beetles this is part of parental care. Larvae feed on vertebrate carcasses which their parents smear with exudates that inhibit microbial growth. We have sequenced the transcriptome of the burying beetle *Nicrophorus vespilloides* and identified six genes that encode lysozymes – a type of antimicrobial enzyme that has previously been implicated in social immunity in burying beetles. When females start breeding and producing antimicrobial anal exudates, we found that the expression of one of these genes was increased by ∼1000 times to become one of the most abundant transcripts in the transcriptome. We conclude that we have likely identified a gene for social immunity, and that it was recruited during evolution from a previous function in personal immunity.

## Introduction

Insects occupy some of the most microbe-rich environments in nature and have evolved diverse immunological defences to overcome the challenge that microbes pose to their fitness [1,2]. In some group-living species, individuals are selected to defend other individuals, as well as themselves, from potential pathogens. This is social immunity in the broad sense, and it is seen in transient animal societies such as animal families as well as more permanent animal societies such as the eusocial insects and group-living primates [3]. Social immunity can take a range of forms, from the collective behaviour that causes social fever in bees, to the production of antibacterial substances by parents to defend offspring or a breeding resource [2–4]. Yet, while the mechanisms underlying personal immunity in insects are increasingly well-described [5,6], relatively little is known about the mechanisms underlying social immunity. Nor is it clear whether social immune function might have originally been derived from personal immune function.

In burying beetles (*Nicrophorus* spp), social immunity is a vital part of parental care. These insects breed on small vertebrate carcasses which they shave, roll into a ball and smear with anal exudates. These exudates have strong antimicrobial properties [7,8] and promote larval survival [9]. The strength of antimicrobial activity in anal exudates is proportional to the perceived microbial threat but increasing levels of antimicrobial activity comes at a fitness cost to adults [10] and trades-off against personal immunity [11]. Antimicrobial activity in the anal exudates is thus carefully modulated. It is virtually non-existent in non-breeding individuals [8], is induced by reproduction and the presentation of a carcass [8] and reaches its maximum strength when the larvae arrive at the carcass shortly after hatching in the soil surrounding the carcass [11].

How has social immunity evolved in the burying beetle? One hypothesis is that elements of the personal immune response have been recruited to control the microbiota in the wider environment. Lysozymes, which are enzymes that can kill bacteria by hydrolysing structural polysaccharides in their cell walls, are a likely candidate because they are ubiquitous in nature and have key roles in personal immunity [5,12]. In insects that feed on microbe-rich resources (e.g. *Drosophila*, house-fly), lysozymes in the gut are thought not only to have an immune function but also to digest bacteria [13,14]. Perhaps in burying beetles, lysozymes that were originally confined to the gut are now exuded and applied to the carcass to limit microbial growth during reproduction. Supporting this hypothesis is the finding that a key active antimicrobial substance in the burying beetle’s anal exudates has lysozyme-like properties [8,9]

Here our aim is to test whether lysozyme genes are upregulated during the mounting of a social immune response in the burying beetle *N. vespilloides*. We sequence the *N. vespilloides* transcriptome, identify the lysozymes within it, and compare the transcriptional response in the gut of breeding and non-breeding female burying beetles to identify upregulated genes.

### Methods

#### Beetle rearing and dissecting

The beetles used in this experiment were bred in 2014 and descended from field-collected beetles trapped earlier that year from two sites in Cambridgeshire, UK. The field-collected beetles were interbred to create a large, genetically diverse population. This population was maintained with full parental care and no inbreeding for several generations before the start of this experiment.

We examined the transcriptional response to breeding in *N. vespilloides* by comparing the transcriptional profiles of a breeding female beetle and a non-breeding female of the same age. We focused on females alone, because our previous work suggests that they contribute more to social immunity than males [1]. Prior to each treatment, beetles were given a small meal of minced beef as part of the usual protocol for beetle husbandry in the lab. The “breeding” treatment initially consisted of 3 female-male pairs of beetles. Each pair was placed in a breeding box with soil and a thawed mouse carcass (10-16 g). These boxes were then put in a dark cupboard to simulate underground conditions and the beetles were allowed to mate and begin preparing the carcass. Forty-eight hours after pairing, at peak antimicrobial activity in the anal exudates [2], we removed the female from each breeding box and placed them individually in small plastic boxes (box dimensions, length x width x depth: 12 cm x 8 cm x 2 cm) where they remained for approximately 1 hour before being killed and dissected. The “non-breeding” treatment consisted of 3 females that were treated in exactly the same way as the “breeding” treatment except that the non-breeding females were placed individually in a breeding box that did not contain a mouse carcass. This was repeated on two occasions to generate 6 breeding and 6 non-breeding beetles. Individual beetles were euthanized with CO_2_ and immediately dissected to remove the gut. We focused on gut tissue because this is where the anal exudates are produced.

#### Transcriptome Sequencing

The dissected gut was immediately homogenised in TRIzol® Reagent, (Life Technologies) and frozen in liquid nitrogen. RNA was extracted following the standard protocol. Illumina sequencing libraries were constructed with poly-A enrichment and sequenced in a single lane of Illumina HiSeq 2500® (version 3 chemistry, 100bp paired-end reads) by BGI (Hong Kong). Raw reads were initially checked for quality using FastQC [3]. Having been found to be satisfactory, they were then trimmed using Trimmomatic [4], removing trailing and leading bases with a quality below q15, cutting reads where quality fell below q20 in a 4 base sliding window, and only retaining reads of minimum length 30. The raw sequence data has been submitted to the NCBI sequence read archive under the accession number xxxxxxxxx.

#### Transcriptome assembly

The RNAseq reads from a single breeding and a single non-breeding beetle gut were combined and the transcriptome *de novo* assembled (Supplementary Figure S1). The assembly was performed using Trinity, a compact and fast transcript assembly program for Illumina RNA-seq data [5]. Briefly, a single Trinity assembly was built using forward and reverse reads from both libraries and default parameters. The full recommended protocol “Identification and Analysis of Differentially Expressed Trinity Genes and Transcripts” was applied (http://trinityrnaseq.sourceforge.net/analysis/diff_expression_analysis.html, accessed 10/04/15). The assembled transcriptome has been submitted to the NCBI short read archive (accession xxxxxxxxxxxx) and predicted peptides are available in Dryad xxxxxxxxxx.

#### Differential expression analysis

To estimate transcript abundance, we aligned reads separately from each library onto the combined*-*read transcriptome assembly using the short read aligner bowtie [6]. Abundance estimates were then produced using RSEM [7]. These steps are combined into a single perl script bundled with Trinity, 

~~~
align_and_estimate_abundance.pl
~~~

. In further analyses we used estimates of transcript abundance for each gene (as opposed to each isoform). Finally, we estimated levels of differential expression using EdgeR, an R Bioconductor package for differential expression analysis. Differentially expressed transcripts were identified using the Trinity scripts 

~~~
run_DE_analysis.pl
~~~

 and 

~~~
analyze_diff_expr.pl
~~~

 with default settings. In a very small number of cases it was clear that alternative haplotypes of a gene had been split into two genes during the assembly and this gave a false signal of differential expression. The avoid this we identified all the genes whose predicted peptides were >98% identical to another gene using CD-HIT [8] and excluded them from the analysis.

#### Peptide and domain prediction

Trinity assembles nucleotide reads into nucleotide transcripts, and as such candidate peptide sequences must be predicted post-hoc (Supplementary Figure S1). Peptide predictions were generated from the combined read assembly using Transdecoder [5] and the standard protocol for peptide prediction. Any transcript that did not encode a predicted peptide was removed from our assembly.

The resulting peptide predictions were then run through the NCBI Batch Conserved Domain Search [9] to annotate domains. Putative lysozymes were then identified by presence of the LYZ1 C-type lysozyme domain (cd00119), which is found in *Drosophila* lysozymes and expected to be required for *Drosophila-*like function of the protein. Full domain annotations can be found in the Dryad xxxxxxxxxx. The predicted lysozyme sequences have been submitted to genbank under the accession numbers xxxxxxxxxxxxx.

#### Quantitative PCR

Differential expression of lysozymes was verified by quantitative PCR. We synthesised cDNA using Promega Go-script® reverse transcriptase following the standard conditions using 1ul RNA template and incubating at 25C for 5 minutes, 50C for 50 minutes and 70C for 15 minutes. PCR primers were designed that amplify the six lysozymes (all three *lys1* isoforms were amplified by a single primer pair) and the reference gene *actin5C* (Table S1). The quantitative PCR was performed using SYBR green using the SensiFAST SYBR Hi-ROX Kit with a 10 ul reaction volume (2 ul template cDNA diluted 1:10 from original cDNA synthesis).

### Results

#### The burying beetle transcriptome

To allow us to investigate the transcriptional response in the guts of burying beetle when they breed, we first sequenced their transcriptome from the gut of a single breeding and non-breeding beetle, combined the sequence reads and then assembled them *de novo.* This process resulted in 11290 genes that encoded 26378 different transcripts. This suggests that we sequenced the majority of genes in the genome, as the exceptionally well-annotated *Drosophila* genome contains 13920 protein coding genes encoding 30443 transcripts (Flybase release 6). As the guts we used for the RNA extraction might contain RNA from the mouse the beetles were feeding on, or nematode parasites, we used Blast to search for the most similar sequence in the *Mus musculus, C. elegans, Drosophila melanogaster* and *Tribolium castaneum* genomes. The top hit of 91% of the genes was another insect (*Drosophila* or the beetle *Tribolium*), suggesting the levels of contamination were low (Figure 1A).

**Figure 1.**
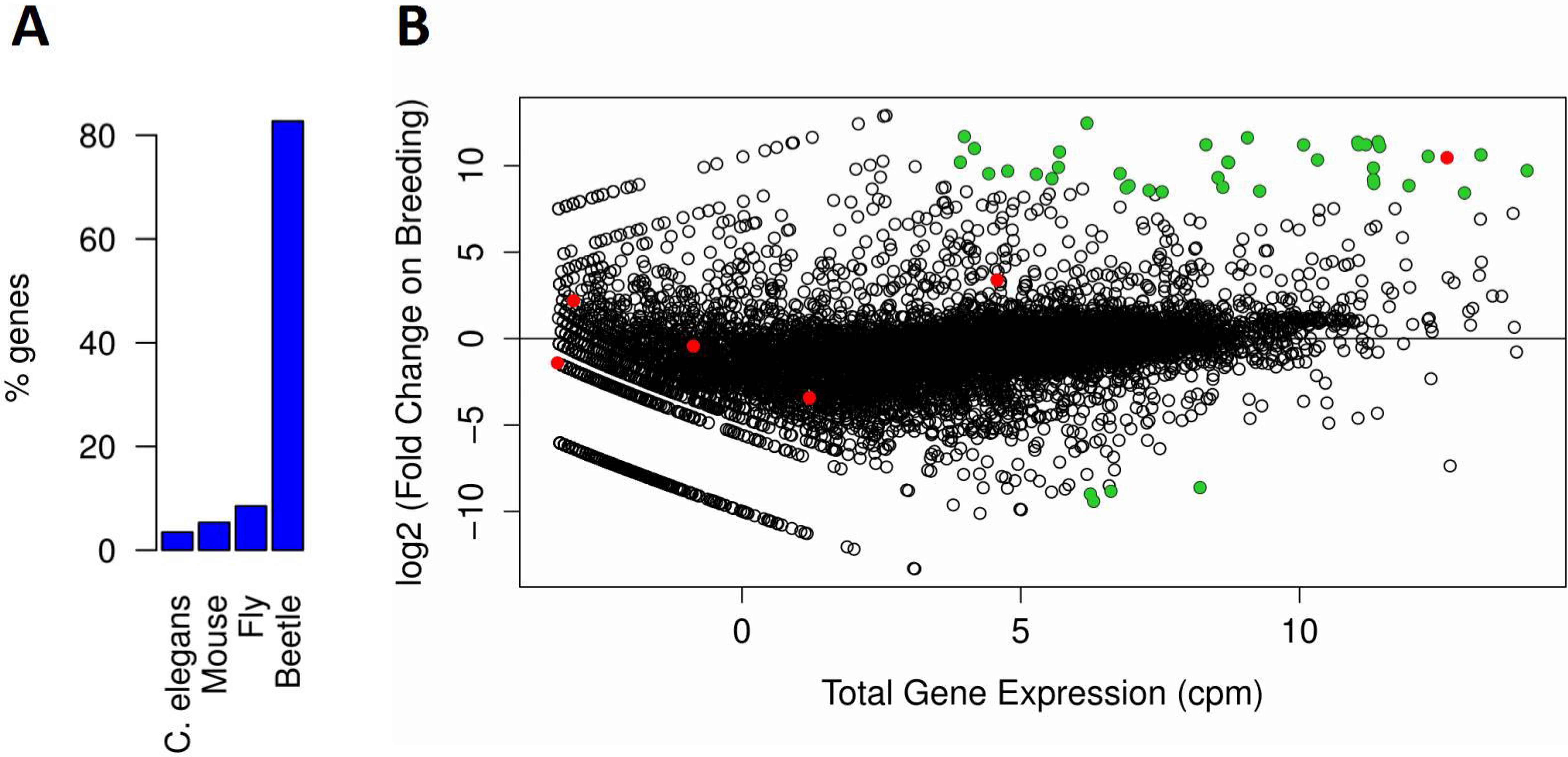
The transcriptome of *N. vespilloides.* (A) The percentage of peptides whose most similar sequence was in the genome of the mouse *Mus musculus*, the nematode *Caenorhabditis elegans*, the fly *Drosophila melanogaster* or the beetle *Tribolium castaneum.* A single isoform of each gene in the differential expression analysis is included. (B) Total gene expression (counts per million) and log_2_ (fold change) in gene expression in the guts of breeding versus non-breeding females. Lysozymes are shown in red. The most significantly differentially expressed genes (*P*<10^-20^) are in green.

#### Many genes are strongly upregulated in the guts of breeding beetles

By mapping reads from the breeding and non-breeding beetles to the transcriptome, we found that there was a strong transcriptional response in the breeding beetles (Figure 1B). Among the most significantly differentially expressed genes (*p*<10^20^), 90% were upregulated in the breeding beetles (Figure 1B; *N*=42, 95% binomial CI: 77-97%). The magnitude of these changes in transcription was often large – on average the expression of these 42 most significantly differentially expressed genes changed by nearly 1000 times (mean log_2_ (fold change)=9.96). Furthermore, some of the most strongly differentially expressed genes also had the highest total levels of expression in our transcriptome (Figure 1B).

#### A lysozyme is highly expressed in breeding females

We identified lysozymes by searching for the conserved LYZ1 domain, which contains the active site of C-type lysozymes. Using this approach we identified six lysozymes (Figure 2A). These ranged in size from 103-214 amino acids, which is within the typical size range of insect lysozymes.

**Figure 2.**
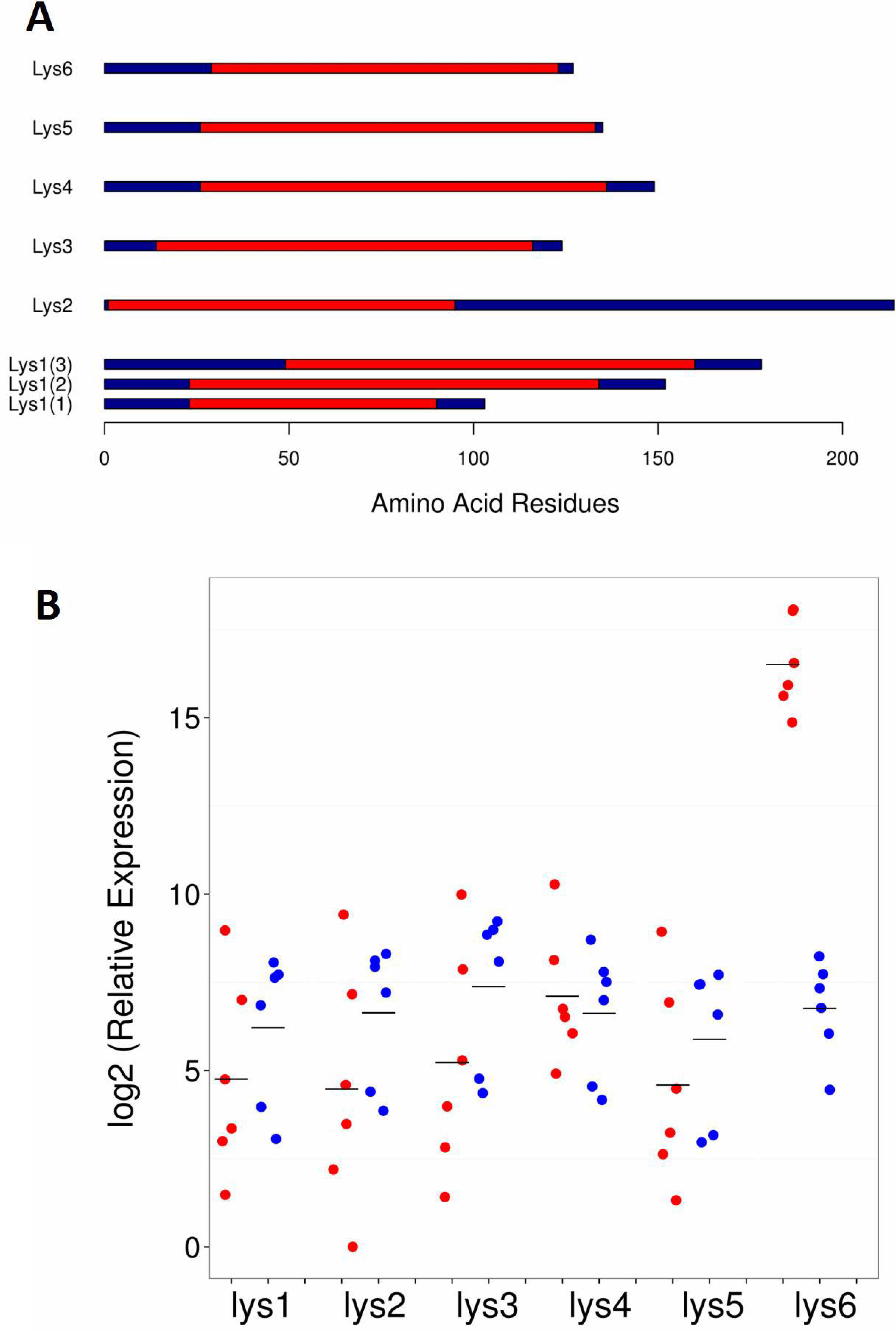
Lysozymes and their expression. (A) The six predicted lysozymes in the transcriptome of *N. vespilloides.* The LYZ1 C-type lysozyme domain (cd00119) is shown in red. There are three alternative isoforms of *Lys1.* (B) The expression of the lysozyme genes in the guts of six breeding (red) and six non-breeding (blue) females. Expression was measured by quantitative PCR relative to *Actin5C* (scale shifted so begin at zero). Each point is the mean of three technical replicates and the horizontal bars are means.

To identify the gene that may be responsible for the antimicrobial activity of the anal exudate of breeding females we compared the expression of the six lysozyme genes in breeding and non-breeding females in the whole transcriptome data. Five of the genes had similar expression levels in breeding and non-breeding beetles, while *Lys6* was massively upregulated – the expression level in breeding females was 1409 times greater than in non-breeding females (Figure 1B; log_2_ (fold change)=10.46, *p*<10^-26^). In the breeding beetles *Lys6* was the 14^th^ most abundant transcript in the entire transcriptome, while in the non-breeding beetles it was only the 5967^th^ most abundant.

We replicated this result using quantitative PCR to measure the expression of the lysozymes across six breeding and six non-breeding females (Figure 2B). In the non-breeding females the different lysozymes all had similar levels of expression. As was the case in the transcriptome analysis, *Lys6* was strongly upregulated in breeding females, with an average expression level that was 860 times than non-breeding beetles (*t* =12.6, df = 9.99, *p* <10^-07^). The expression of the five remaining lysozymes was unaltered in the breeding females.

### Discussion

Our analyses indicate that breeding induces a very strong transcriptional response in female burying beetles, causing substantial upregulation of just one lysozyme gene (*Lys6*) in their gut tissues relative to non-breeding females. This correlates with our earlier phenotypic observations that the antimicrobial properties of the anal exudates in females are dramatically elevated after the presentation of a carcass and the onset of reproduction [8, 11]. Since none of the other 5 lysozyme genes that are expressed in the gut were upregulated during reproduction, it suggests that upregulation of *Lys6* causes at least some of the change in the exudates’ antimicrobial properties during breeding. In addition, it provides further evidence to support the hypothesis that burying beetles have recruited a component of their personal immune system, namely lysozyme, to play a major role in social immunity.

Does this mean we have therefore found a gene for social immunity in the burying beetle *N. vespilloides*? We cautiously suggest that the answer to this question is likely to be yes. Previous work has identified lysozyme-like activity in the antimicrobial anal exudates [8,9]. Given the very high levels of *Lys6* expression were only detected in breeding females, it seems most likely that this gene functions primarily in social immunity. The finding that a lysozyme has a role in social immunity would be unsurprising. These enzymes provide a broad-spectrum defence against microbes in their environment, and they are secreted onto external surfaces that are vulnerable to infection, such as the gut, eyes, mucous membranes and respiratory tract [12]. It may therefore be straightforward to recruit lysozymes to social immune functions. Nevertheless, further experiments are now needed to confirm that the protein encoded by this gene is present in the anal exudates and that it is indeed functioning in social immunity.

It might be argued that bacteria form a key part of the diet of breeding burying beetles or their larvae, but not of non-breeding burying beetles. Thus, a possible alternative interpretation of our data is that *Lys6* primarily serves a digestive function, rather than an immune function, as has been suggested for the lysozymes expressed in housefly or *Drosophila* guts. However, we think this alternative interpretation is unlikely as behavioural evidence suggests that beetles prefer to feed on meat rather than on the microbes living on the meat [7]. Furthermore, beetles in both treatments were given a similar meat-based diet, whether they bred or not, which suggests that upregulation of *Lys6* in the breeding beetles was not induced simply by consuming mouse flesh. Thus, although at this stage we cannot rule out the possibility that *Lys6* plays some minor role in digestion, this is unlikely to be its sole or even primary function.

In summary, we suggest that we have found a gene (*Lys 6*) for social immunity in the burying beetle and that it was recruited from personal immune function in the evolutionary past. The challenge for future work is to determine how this gene’s function is integrated with other components of the social immune system to influence the microbial community on the burying beetle’s carcass breeding resource.

## Acknowledgements

This work was funded by ERC grant DrosophilaInfection 281668 to FMJ, NERC grant NE/H019731/1 to RMK, ERC grant BALDWINIAN_BEETLES 310785 to RMK and a Wolfson Merit Award to RMK. We thank S. Aspinall for help in maintaining the burying beetle population.

